## Supplemental Figures and Tables for "Spinal V1 inhibitory interneuron clades differ in birthdate, projections to motoneurons, and heterogeneity"

**Abbreviated title**: Diversity of V1 Spinal Interneurons

**Authors**

Andrew E. Worthy[1,2], JoAnna T. Anderson[2], Alicia R. Lane[2], Laura Gomez-Perez[2], Anthony A. Wang[1], Ronald W. Griffith[1,2], Andre F. Rivard[2], Jay B. Bikoff[3], Francisco J. Alvarez[1,2#]

[1] Department of Physiology, Emory University School of Medicine, Atlanta, GA, USA 30322

[2] Department of Cell Biology, Emory University School of Medicine, Atlanta, GA, USA 30322

[3] Department of Developmental Neurobiology, St Jude Children's' Research Hospital, Memphis, TN, USA 38105

**Supplementary Table S1;** related to Figure 4D.

| *Ordinary one-way ANOVA summary* | |
|---|---|
| *F* | 43.30 |
| *P value* | <0.0001 |
| *P value summary* | **** |
| *Significant diff. among means (P < 0.05)?* | Yes |
| *R squared* | 0.9319 |

| *Bonferroni's multiple comparisons test* | *Mean Diff.* | *95.00% CI of diff.* | *t (df:19)* | *Summary* | *Adjusted P Value* |
|---|---|---|---|---|---|
| *Th13 vs. L1* | -11.39 | -26.33 to 3.552 | 2.670 | ns | 0.3179 |
| *Th13 vs. L2* | -14.72 | -28.70 to -0.7460 | 3.690 | * | 0.0327 |
| *Th13 vs. L3* | -27.47 | -41.45 to -13.50 | 6.885 | **** | <0.0001 |
| *Th13 vs. L4/5* | -41.50 | -55.48 to -27.52 | 10.40 | **** | <0.0001 |
| *Th13 vs. L6* | -42.07 | -55.43 to -28.70 | 11.03 | **** | <0.0001 |
| *Th13 vs. S1* | -0.111 | -15.05 to 14.83 | 0.026 | ns | >0.9999 |
| *L1 vs. L2* | -3.333 | -17.31 to 10.64 | 0.835 | ns | >0.9999 |
| *L1 vs. L3* | -16.08 | -30.06 to -2.107 | 4.031 | * | 0.0150 |
| *L1 vs. L4/5* | -30.11 | -44.09 to -16.13 | 7.546 | **** | <0.0001 |
| *L1 vs. L6* | -30.68 | -44.04 to -17.31 | 8.041 | **** | <0.0001 |
| *L1 vs. S1* | 11.28 | -3.663 to 26.22 | 2.644 | ns | 0.3362 |
| *L2 vs. L3* | -12.75 | -25.69 to 0.1895 | 3.451 | ns | 0.0562 |
| *L2 vs. L4/5* | -26.78 | -39.72 to -13.84 | 7.249 | **** | <0.0001 |
| *L2 vs. L6* | -27.34 | -39.62 to -15.07 | 7.802 | **** | <0.0001 |
| *L2 vs. S1* | 14.61 | 0.6349 to 28.59 | 3.662 | * | 0.0348 |
| *L3 vs. L4/5* | -14.03 | -26.97 to -1.088 | 3.797 | * | 0.0256 |
| *L3 vs. L6* | -14.59 | -26.87 to -2.319 | 4.164 | * | 0.0111 |
| *L3 vs. S1* | 27.36 | 13.38 to 41.34 | 6.857 | **** | <0.0001 |
| *L4/5 vs. L6* | -0.567 | -12.84 to 11.71 | 0.162 | ns | >0.9999 |
| *L4/5 vs. S1* | 41.39 | 27.41 to 55.37 | 10.37 | **** | <0.0001 |
| *L6 vs. S1* | 41.96 | 28.59 to 55.32 | 11.00 | **** | <0.0001 |

**Supplementary Table S2;** related to Figure 5D.

Nested One-Way ANOVA $F_{(11, 80)} = 36.09$; $p < 0.0001$

Variation within animals: SD = 3.018, Variance = 9.108

Inter-animal variation for each motor column/segment: Chi square$_{(1)}$ = 0.5061; p = 0.4768

| *Bonferroni's multiple comparisons test* | *Mean Diff.* | *95.00% CI of diff.* | *t (df:80)* | *Summary* | *Adjusted P Value* |
|---|---|---|---|---|---|
| *Th12/Th13 HMC vs. L1/L2 LMCv* | 0.7566 | -2.819 to 4.332 | 0.741 | ns | >0.9999 |
| *Th12/Th13 HMC vs. L4/L5 LMCd* | -3.142 | -6.515 to 0.2315 | 3.262 | ns | 0.1074 |
| *Th12/Th13 HMC vs. L4/L5 LMCv* | -5.112 | -8.384 to -1.839 | 5.471 | **** | <0.0001 |
| *Th12/Th13 HMC vs. L6 LMCd* | 0.8142 | -2.697 to 4.325 | 0.812 | ns | >0.9999 |
| *Th12/Th13 HMC vs. Th12/Th13 MMC* | 2.695 | -0.8050 to 6.195 | 2.697 | ns | 0.5632 |
| *Th12/Th13 HMC vs. L1/L2 MMC* | 2.551 | -0.9381 to 6.040 | 2.561 | ns | 0.8136 |
| *Th12/Th13 HMC vs. L3/L4 MMC* | -0.3096 | -3.682 to 3.062 | 0.321 | ns | >0.9999 |
| *Th12/Th13 HMC vs. S1/2 MMC* | 2.373 | -0.9170 to 5.663 | 2.526 | ns | 0.8913 |
| *Th12/Th13 HMC vs. Th12/Th13 PGC* | 7.144 | 3.876 to 10.41 | 7.658 | **** | <0.0001 |
| *Th12/Th13 HMC vs. L1/L2 PGC* | 7.097 | 3.642 to 10.55 | 7.195 | **** | <0.0001 |
| *Th12/Th13 HMC vs. S1/S2 PGC* | 5.473 | 2.110 to 8.836 | 5.700 | **** | <0.0001 |
| *L1/L2 LMCv vs. L4/L5 LMCd* | -3.898 | -7.275 to -0.5210 | 4.043 | ** | 0.0080 |
| *L1/L2 LMCv vs. L4/L5 LMCv* | -5.868 | -9.145 to -2.592 | 6.273 | **** | <0.0001 |
| *L1/L2 LMCv vs. L6 LMCd* | 0.05760 | -3.457 to 3.572 | 0.057 | ns | >0.9999 |
| *L1/L2 LMCv vs. Th12/Th13 MMC* | 1.938 | -1.565 to 5.442 | 1.938 | ns | >0.9999 |
| *L1/L2 LMCv vs. L1/L2 MMC* | 1.794 | -1.699 to 5.287 | 1.799 | ns | >0.9999 |
| *L1/L2 LMCv vs. L3/L4 MMC* | -1.066 | -4.442 to 2.310 | 1.106 | ns | >0.9999 |
| *L1/L2 LMCv vs. S1/2 MMC* | 1.616 | -1.678 to 4.911 | 1.719 | ns | >0.9999 |
| *L1/L2 LMCv vs. Th12/Th13 PGC* | 6.387 | 3.116 to 9.659 | 6.838 | **** | <0.0001 |
| *L1/L2 LMCv vs. L1/L2 PGC* | 6.341 | 2.882 to 9.800 | 6.421 | **** | <0.0001 |
| *L1/L2 LMCv vs. S1/S2 PGC* | 4.716 | 1.349 to 8.084 | 4.906 | *** | 0.0003 |
| *L4/L5 LMCd vs. L4/L5 LMCv* | -1.970 | -5.024 to 1.084 | 2.259 | ns | >0.9999 |
| *L4/L5 LMCd vs. L6 LMCd* | 3.956 | 0.6471 to 7.264 | 4.187 | ** | 0.0048 |
| *L4/L5 LMCd vs. Th12/Th13 MMC* | 5.836 | 2.540 to 9.133 | 6.200 | **** | <0.0001 |
| *L4/L5 LMCd vs. L1/L2 MMC* | 5.692 | 2.407 to 8.977 | 6.069 | **** | <0.0001 |
| *L4/L5 LMCd vs. L3/L4 MMC* | 2.832 | -0.3288 to 5.993 | 3.138 | ns | 0.1572 |
| *L4/L5 LMCd vs. S1/2 MMC* | 5.514 | 2.441 to 8.588 | 6.285 | **** | <0.0001 |
| *L4/L5 LMCd vs. Th12/Th13 PGC* | 10.29 | 7.236 to 13.33 | 11.82 | **** | <0.0001 |
| *L4/L5 LMCd vs. L1/L2 PGC* | 10.24 | 6.990 to 13.49 | 11.04 | **** | <0.0001 |
| *L4/L5 LMCd vs. S1/S2 PGC* | 8.615 | 5.463 to 11.77 | 9.575 | **** | <0.0001 |
| *L4/L5 LMCv vs. L6 LMCd* | 5.926 | 2.720 to 9.132 | 6.474 | **** | <0.0001 |
| *L4/L5 LMCv vs. Th12/Th13 MMC* | 7.807 | 4.613 to 11.00 | 8.561 | **** | <0.0001 |
| *L4/L5 LMCv vs. L1/L2 MMC* | 7.663 | 4.481 to 10.84 | 8.435 | **** | <0.0001 |
| *L4/L5 LMCv vs. L3/L4 MMC* | 4.802 | 1.749 to 7.855 | 5.509 | **** | <0.0001 |
| *L4/L5 LMCv vs. S1/2 MMC* | 7.485 | 4.523 to 10.45 | 8.850 | **** | <0.0001 |
| *L4/L5 LMCv vs. Th12/Th13 PGC* | 12.26 | 9.319 to 15.19 | 14.62 | **** | <0.0001 |
| *L4/L5 LMCv vs. L1/L2 PGC* | 12.21 | 9.065 to 15.35 | 13.60 | **** | <0.0001 |
| *L4/L5 LMCv vs. S1/S2 PGC* | 10.58 | 7.542 to 13.63 | 12.18 | **** | <0.0001 |
| *L6 LMCd vs. Th12/Th13 MMC* | 1.881 | -1.557 to 5.318 | 1.916 | ns | >0.9999 |
| *L6 LMCd vs. L1/L2 MMC* | 1.737 | -1.690 to 5.163 | 1.775 | ns | >0.9999 |
| *L6 LMCd vs. L3/L4 MMC* | -1.124 | -4.431 to 2.184 | 1.190 | ns | >0.9999 |
| *L6 LMCd vs. S1/2 MMC* | 1.559 | -1.665 to 4.783 | 1.694 | ns | >0.9999 |
| *L6 LMCd vs. Th12/Th13 PGC* | 6.330 | 3.129 to 9.530 | 6.926 | **** | <0.0001 |
| *L6 LMCd vs. L1/L2 PGC* | 6.283 | 2.891 to 9.675 | 6.488 | **** | <0.0001 |
| *L6 LMCd vs. S1/S2 PGC* | 4.659 | 1.361 to 7.957 | 4.947 | *** | 0.0003 |
| *Th12/Th13 MMC vs. L1/L2 MMC* | -0.1440 | -3.559 to 3.271 | 0.148 | ns | >0.9999 |

| | | | | | |
|---|---|---|---|---|---|
| *Th12/Th13 MMC vs. L3/L4 MMC* | -3.004 | -6.300 to 0.2912 | 3.193 | ns | 0.1330 |
| *Th12/Th13 MMC vs. S1/2 MMC* | -0.3218 | -3.533 to 2.890 | 0.351 | ns | >0.9999 |
| *Th12/Th13 MMC vs. Th12/Th13 PGC* | 4.449 | 1.261 to 7.637 | 4.887 | *** | 0.0003 |
| *Th12/Th13 MMC vs. L1/L2 PGC* | 4.403 | 1.022 to 7.783 | 4.562 | ** | 0.0012 |
| *Th12/Th13 MMC vs. S1/S2 PGC* | 2.778 | -0.5082 to 6.065 | 2.961 | ns | 0.2664 |
| *L1/L2 MMC vs. L3/L4 MMC* | -2.860 | -6.144 to 0.4236 | 3.051 | ns | 0.2044 |
| *L1/L2 MMC vs. S1/2 MMC* | -0.1778 | -3.378 to 3.022 | 0.195 | ns | >0.9999 |
| *L1/L2 MMC vs. Th12/Th13 PGC* | 4.593 | 1.417 to 7.769 | 5.064 | *** | 0.0002 |
| *L1/L2 MMC vs. L1/L2 PGC* | 4.547 | 1.178 to 7.916 | 4.727 | *** | 0.0006 |
| *L1/L2 MMC vs. S1/S2 PGC* | 2.922 | -0.3526 to 6.197 | 3.125 | ns | 0.1633 |
| *L3/L4 MMC vs. S1/2 MMC* | 2.683 | -0.3893 to 5.754 | 3.059 | ns | 0.1997 |
| *L3/L4 MMC vs. Th12/Th13 PGC* | 7.453 | 4.406 to 10.50 | 8.566 | **** | <0.0001 |
| *L3/L4 MMC vs. L1/L2 PGC* | 7.407 | 4.159 to 10.65 | 7.987 | **** | <0.0001 |
| *L3/L4 MMC vs. S1/S2 PGC* | 5.783 | 2.633 to 8.933 | 6.430 | **** | <0.0001 |
| *S1/2 MMC vs. Th12/Th13 PGC* | 4.771 | 1.814 to 7.727 | 5.652 | **** | <0.0001 |
| *S1/2 MMC vs. L1/L2 PGC* | 4.724 | 1.562 to 7.887 | 5.232 | **** | <0.0001 |
| *S1/2 MMC vs. S1/S2 PGC* | 3.100 | 0.03801 to 6.162 | 3.546 | * | 0.0434 |
| *Th12/Th13 PGC vs. L1/L2 PGC* | -0.04630 | -3.185 to 3.093 | 0.052 | ns | >0.9999 |
| *Th12/Th13 PGC vs. S1/S2 PGC* | -1.671 | -4.708 to 1.367 | 1.926 | ns | >0.9999 |
| *L1/L2 PGC vs. S1/S2 PGC* | -1.624 | -4.863 to 1.614 | 1.757 | ns | >0.9999 |

.

**Supplementary Table S3**; related to Figure 5E, top graph.

**Foxp2-V1 vs non-Foxp2-V1 synapses**

Two-Way ANOVA for Synapse Origin and Motor Column/Segment

- Column/Segment: $F_{(11, 248)} = 22.52$, $p < 0.0001$
- Foxp2 vs Non-Foxp2: $F_{(1, 248)} = 6.607$, $p = 0.0107$
- Interaction: $F_{(11, 248)} = 10.99$, $p < 0.0001$

| *Bonferroni's multiple comparisons test* | *Predicted (LS) mean diff.* | *95.00% CI of diff.* | *t (df:248)* | *Summary* | *Adjusted P Value* |
|---|---|---|---|---|---|
| *HMC Th13* | 3.218 | 0.2757 to 6.161 | 3.163 | * | 0.0211 |
| *LMCv L1/L2* | -0.03900 | -2.982 to 2.904 | 0.038 | ns | >0.9999 |
| *LMCd L4/L5* | 4.704 | 2.497 to 6.911 | 6.164 | **** | <0.0001 |
| *LMCv L4/L5* | 4.028 | 1.748 to 6.307 | 5.110 | **** | <0.0001 |
| *LMCd L6* | 4.145 | 0.5405 to 7.749 | 3.326 | * | 0.0122 |
| *MMC Th13* | 0.8494 | -1.942 to 3.641 | 0.878 | ns | >0.9999 |
| *MMC L1/L2* | -0.9560 | -4.077 to 2.165 | 0.886 | ns | >0.9999 |
| *MMC L3/L4* | -0.1519 | -2.556 to 2.252 | 0.183 | ns | >0.9999 |
| *MMC S1* | -4.833 | -7.152 to -2.513 | 6.025 | **** | <0.0001 |
| *PGC Th13* | -0.4782 | -2.927 to 1.970 | 0.565 | ns | >0.9999 |
| *PGC L1/L2* | -0.4767 | -3.268 to 2.315 | 0.494 | ns | >0.9999 |
| *PGC S1* | -1.608 | -4.157 to 0.9403 | 1.825 | ns | 0.8305 |

.

**Supplementary Table S4;** related to Figure 5E, bottom graph.

**Foxp2-V1 vs CB+-V1 (Renshaw) synapses**

Two-Way ANOVA for Synapse origin and Motor Column/Segment

- Column/Segment: $F_{(11, 271)} = 27.67$, $p < 0.0001$
- Foxp2 vs CB+(Renshaw): $F_{(1, 271)} = 77.60$, $p < 0.0001$
- Interaction: $F_{(11, 248)} = 14.71$, $p < 0.0001$

| *Bonferroni's multiple comparisons test* | *Predicted (LS) mean diff.* | *95.00% CI of diff.* | *t (df:248)* | *Summary* | *Adjusted P Value* |
|---|---|---|---|---|---|
| *HMC Th13* | 4.348 | 1.971 to 6.725 | 5.286 | **** | <0.0001 |
| *LMCv L1/L2* | 0.4498 | -1.927 to 2.827 | 0.547 | ns | >0.9999 |
| *LMCd L4/L5* | 5.128 | 2.996 to 7.260 | 6.951 | **** | <0.0001 |
| *LMCv L4/L5* | 6.682 | 4.751 to 8.613 | 9.999 | **** | <0.0001 |
| *LMCd L6* | 5.632 | 3.078 to 8.187 | 6.371 | **** | <0.0001 |
| *MMC Th13* | 0.8168 | -1.448 to 3.081 | 1.042 | ns | >0.9999 |
| *MMC L1/L2* | -0.7661 | -3.110 to 1.578 | 0.945 | ns | >0.9999 |
| *MMC L3/L4* | 2.165 | 0.1277 to 4.202 | 3.071 | * | 0.0282 |
| *MMC S1* | -2.069 | -4.068 to -0.0705 | 2.992 | * | 0.0363 |
| *PGC Th13* | 0.2166 | -1.732 to 2.165 | 0.321 | ns | >0.9999 |
| *PGC L1/L2* | 0.5643 | -1.660 to 2.789 | 0.733 | ns | >0.9999 |
| *PGC S1* | 0.03761 | -2.043 to 2.118 | 0.052 | ns | >0.9999 |

**Supplementary Table S5;** related to *Figure 6C (left: OTP/Foxp2).*

| *Ordinary one-way ANOVA summary* | |
|---|---|
| *F* | 45.10 |
| *P value* | <0.0001 |
| *P value summary* | **** |
| *Significant diff. among means (P < 0.05)?* | Yes |
| *R squared* | 0.9442 |

| *Bonferroni's multiple comparisons test* | *Mean Diff.* | *95.00% CI of diff.* | *t(dg:8)* | *Summary* | *Adjusted P Value* |
|---|---|---|---|---|---|
| *None vs. OTP and Foxp2* | -12.45 | -24.20 to -0.6918 | 3.684 | * | 0.0371 |
| *None vs. OTP only* | 16.67 | 4.917 to 28.42 | 4.934 | ** | 0.0069 |
| *None vs. Foxp2 only* | 22.87 | 11.11 to 34.62 | 6.768 | *** | 0.0009 |
| *OTP and Foxp2 vs. OTP only* | 29.12 | 17.36 to 40.87 | 8.618 | *** | 0.0002 |
| *OTP and Foxp2 vs. Foxp2 only* | 35.31 | 23.56 to 47.07 | 10.45 | **** | <0.0001 |
| *OTP only vs. Foxp2 only* | 6.195 | -5.559 to 17.95 | 1.834 | ns | 0.6243 |

**Supplementary Table S6;** related to *Figure 6C (center: Foxp4/Foxp2).*

| *Ordinary one-way ANOVA summary* | |
|---|---|
| *F* | 81.53 |
| *P value* | <0.0001 |
| *P value summary* | **** |
| *Significant diff. among means (P < 0.05)?* | Yes |
| *R squared* | 0.9683 |

| *Bonferroni's multiple comparisons test* | *Mean Diff.* | *95.00% CI of diff.* | *t(dg:8)* | *Summary* | *Adjusted P Value* |
|---|---|---|---|---|---|
| *None vs. Foxp4 and Foxp2* | -12.45 | 23.68 to 47.30 | 10.45 | **** | <0.0001 |
| *None vs. Foxp4 only* | 16.67 | 39.97 to 63.59 | 15.25 | **** | <0.0001 |
| *None vs. Foxp2 only* | 22.87 | 14.11 to 37.73 | 7.635 | *** | 0.0004 |
| *Foxp4 and Foxp2 vs. OTP only* | 29.12 | 4.478 to 28.10 | 4.798 | ** | 0.0082 |
| *Foxp4 and Foxp2 vs. Foxp2 only* | 35.31 | -21.38 to 2.241 | 2.819 | ns | 0.1352 |
| *Foxp4 only vs. Foxp2 only* | 6.195 | -37.67 to -14.05 | 7.617 | *** | 0.0004 |

**Supplementary Table S7;** related to *Figure 6C (right: OTP/Foxp4).*

| *Ordinary one-way ANOVA summary* | |
|---|---|
| *F* | 117.3 |
| *P value* | <0.0001 |
| *P value summary* | **** |
| *Significant diff. among means (P < 0.05)?* | Yes |
| *R squared* | 0.9778 |

| *Bonferroni's multiple comparisons test* | *Mean Diff.* | *95.00% CI of diff.* | *t(dg:8)* | *Summary* | *Adjusted P Value* |
|---|---|---|---|---|---|
| *None vs. OTP and Foxp4* | 26.05 | 17.11 to 35.00 | 10.14 | **** | <0.0001 |
| *None vs. OTP only* | 21.27 | 12.32 to 30.21 | 8.273 | *** | 0.0002 |
| *None vs. Foxp4 only* | 47.99 | 39.04 to 56.93 | 18.67 | **** | <0.0001 |
| *OTP and Foxp4 vs. OTP only* | -4.787 | -13.73 to 4.156 | 1.862 | ns | 0.5976 |
| *OTP and Foxp4 vs. Foxp4 only* | 21.93 | 12.99 to 30.88 | 8.533 | *** | 0.0002 |
| *OTP only vs. Foxp4 only* | 26.72 | 17.78 to 35.66 | 10.40 | **** | <0.0001 |

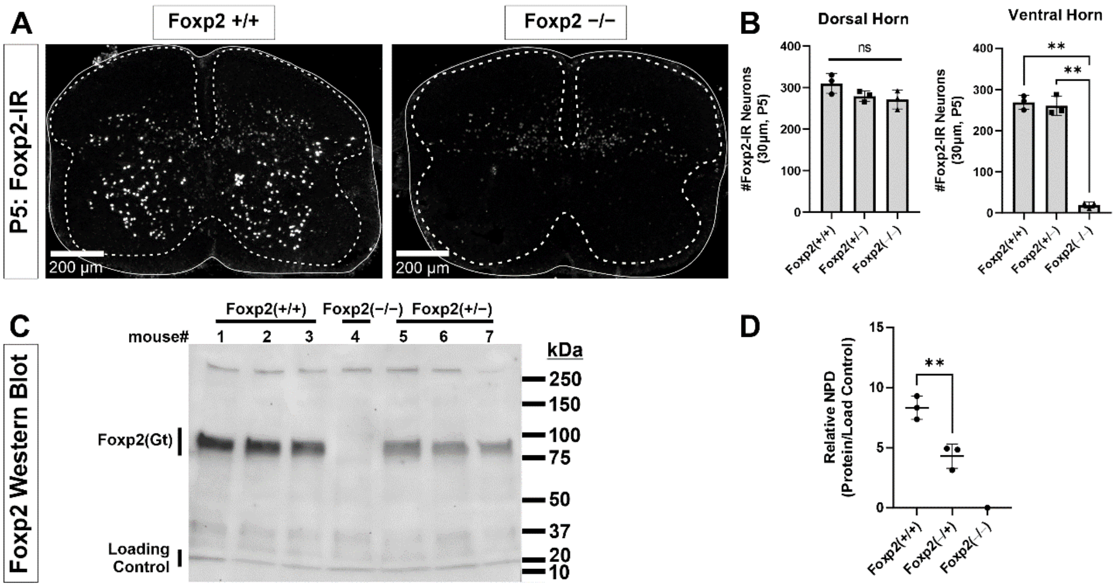


**Supplementary Figure 1: Foxp2 antibody characterization**.

**A,** P5 spinal cords from WT (*Foxp2*$^{+/+}$) and Foxp2 knockout mice (KO: *Foxp2*$^{flpo/flpo}$) immunostained with Santa Cruz goat polyclonal anti-Foxp2 antibody (N-16, Lot#G2911). All nuclear immunostaining in the ventral horn and lateral dorsal horn are absent in KO tissues, however some weak immunoreactivity remains in the dorsal horn. This suggests a weak cross-reaction with another epitope present only in dorsal horn cells.

**B,** Quantification of immunoreactive cell nuclei in the dorsal and ventral horns. For these analyses, a line was drawn horizontally from the top of the central canal to separate dorsal and ventral horns. In 30 µm-thick sections from P5 tissue, the number of Foxp2-IR cells in the dorsal horn is similar in WT (*Foxp2*$^{+/+}$), Foxp2 hets (*Foxp2*$^{flpo/+}$) and Foxp2 KOs (*Foxp2*$^{flpo/flpo}$). Each dot represents one spinal cord serial section from three littermates each with a different genotype. A one-way ANOVA detected no significant differences ($F_{(2,6)} = 3.100$, $p = 0.1190$). This is likely because the number of weakly labeled spots greatly outnumbers the strongly labeled nuclei that disappear from the image. Thus, the comparison does not have enough power for the expected, relatively small difference in number (power for $\alpha = 0.05$ : 0.275 which is smaller than desired power of 0.800). We did not pursue this further because our focus is on ventral horn Foxp2 neurons. In the ventral horn there are mostly strongly labeled nuclei and, correspondingly, all disappear from the spinal cord with only a few exceptions of weak labeling. A one-way ANOVA detected significant differences ($F_{(2,6)} = 3.100$, $p < 0.0001$) and this comparison was adequately powered (power for $\alpha = 0.05$ : 1.000). Post-hoc pair-wise comparisons (Bonferroni corrected t-tests) showed that WT (*Foxp2*$^{+/+}$) and Foxp2 hets (*Foxp2*$^{flpo/+}$) were significantly different to Foxp2 KOs (*Foxp2*$^{flpo/flpo}$) (WT vs KO: $p < 0.0001$ $t(6) = 17.41$; het vs KO: $p < 0.0001$ $t(6) = 16.83$) but not in between them (WT vs het: $p > 0.9999$ $t(6) = 0.5797$) suggesting that heterozygosis did not affect the number of ventral horn interneurons detected using immunolabeling. In summary, our antibody detects Foxp2 expression in the ventral horn with high sensitivity and specificity.

**C,** Antibody specificity confirmed via western blot. Spinal cords from P5 mice from a single litter were extracted and immediately homogenized in a nuclear extraction solution containing protease inhibitors [NE-PER™ Nuclear and Cytoplasmic Extraction (Thermo Fisher); cOmplete™ Protease Inhibitor Cocktail (Roche)]. The nuclear fraction was used for the protein assay and subsequent western blot. Blots were stained with antibodies against Foxp2 (N-16, Santa Cruz) and, as a loading control, against acetyl-Histone H3 (Rabbit, EMD Millipore, 06-599). A single Foxp2 band is revealed at the expected theoretical 80 kDa molecular weight. This band disappears in the Foxp2 KO lane (*Foxp2*$^{flpo/flpo}$).

**D,** Western blot quantification. Protein content was estimated measuring the normalized pixel density (NPD = pixel density in ROI encompassing stained bands – pixel density of same ROI placed just above the band) in the Foxp2 band and relative to the NPD of the respective loading controls in each lane. Foxp2 protein was undetectable in the Foxp2 KO lane (*Foxp2*$^{flpo/flpo}$). In WT animals we detected almost double the amount of Foxp2 protein compared to hets (*Foxp2*$^{flpo/+}$), and this difference was significant (two-tailed t-test $t(4) = 5.004$, $p = 0.0075$). This suggests that Foxp2 gene expression is halved in hets, but this does not affect detectability of Foxp2 ventral horn interneurons which were similar in number in hets vs WTs (see ***B***). In this graph each dot is one animal/lane and the average ±SD indicated.

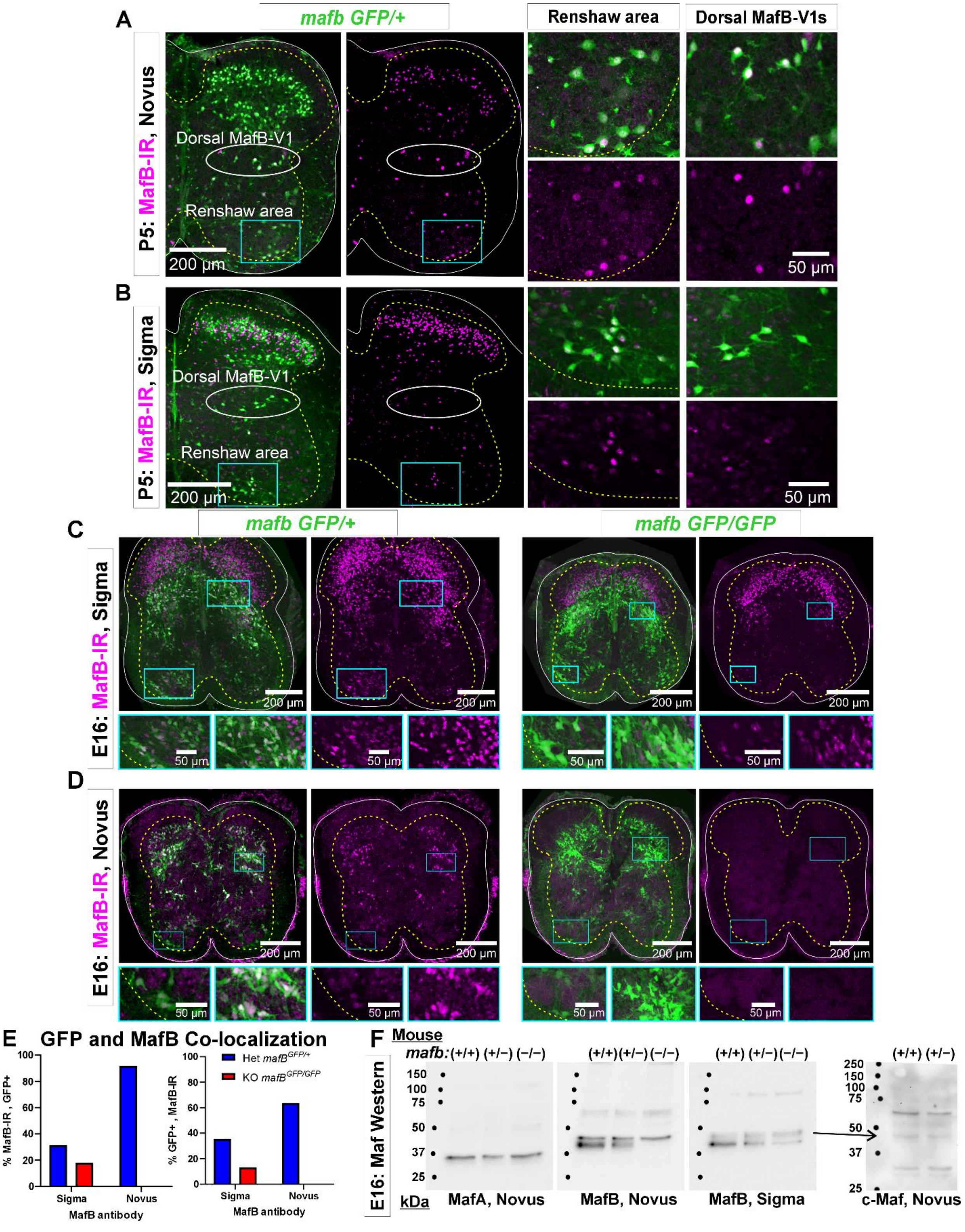

A
mafb GFP/+
Renshaw area
Dorsal MafB-V1s
P5: MafB-IR, Novus
Dorsal MafB-V1
Renshaw area
200 µm
50 µm
B
P5: MafB-IR, Sigma
Dorsal MafB-V1
Renshaw area
200 µm
50 µm
C
mafb GFP/+
mafb GFP/GFP
E16: MafB-IR, Sigma
200 µm
50 µm
D
E16: MafB-IR, Novus
200 µm
50 µm
E
GFP and MafB Co-localization
% MafB-IR , GFP+
% GFP+ , MafB-IR
Het mafB GFP/+
KO mafB GFP/GFP
Sigma
Novus
MafB antibody
F
Mouse
mafb:(+/+) (+/−) (−/−)
E16: Maf Western
kDa
MafA, Novus
MafB, Novus
MafB, Sigma
c-Maf, Novus

**Supplementary Figure 2: Characterization of MafB (Novus, NB600-266) and MafB (Sigma, HPA00563) antibody immunoreactivities in the spinal cord.**

Each antibody was directed against different regions of the mouse MafB protein: aa18-167 for the Sigma antibody (Lot#A31532) and aa100-150 for the Novus antibody (lot#1). These target sequences are shared with mouse MafA and c-Maf, but the overlap is larger with the Sigma antibody immunogen. A BLAST sequence search found 98%, 53% and 64% significant alignment between the MafB sequence used for creation of the Sigma antibody with, respectively, aa18-140 in mouse MafB, aa18-143 in mouse MafA and aa19-118 in mouse c-Maf. The sequence used in the Novus antibody detected no alignment with proteins other than mouse MafB.

**A,** P5 spinal cord from a $mafb^{GFP/+}$ mouse immunostained with MafB-Novus antibodies. GFP (green) reports *mafb* gene expression (see Supplementary Figure 3). Immunohistochemistry detects MafB protein (Cy3, magenta). Higher magnifications of the indicated areas, including the dorsal and ventral (Renshaw) MafB-V1 groups are shown to the right.

**B,** P5 spinal cord from a $mafb^{GFP/+}$ mouse immunostained with MafB-Sigma antibodies. The MafB-Sigma antibody detects many more dorsal horn neurons than the MafB-Novus antibody. Many interneurons at this location are known to express c-Maf (Hu et al., 2012; Frezel et al., 2023). In the ventral horn MafB immunoreactivity is qualitatively similar for both antibodies. This includes both the regions occupied by V1 Renshaw cells and by dorsal-MafB-V1s (Pou6f2-V1 clade).

**C,** Immunoreactivity against MafB-Sigma antibodies in het ($mafb^{GFP/+}$) and KO ($mafb^{GFP/GFP}$) tissue from E16 MafB-GFP reporter mice (the MafB KO is lethal at P0 because it cannot breathe on its own). Low-mag images of MafB-GFP and MafB-immunoreactivity combined (left panel) or MafB-immunoreactivity alone (right panel) in the het (left pair) and the KO mouse (right pair). The boxed ventral and dorsal horn areas are magnified below. GFP in MafB-GFP mice reports activity of the *mafb* promoter, but the knocked-in GFP inactivates the *mafb* allele. When both alleles carry GFP (KO, $mafb^{GFP/GFP}$), GFP reports cells with gene expression from the *mafb* locus although no *mafb* mRNA or protein is produced. In the het mouse one allele produces *mafb* mRNA: in this tissue, there is a high degree of overlap between GFP and protein immunoreactivity in the ventral and deep dorsal horns. However, there is more MafB-Sigma immunoreactivity than GFP in superficial laminae. In the KO, ventral horn immunoreactivity is greatly diminished, but lingering weak immunoreactivity remains in many neurons, including Renshaw cells. This could represent cross-reaction with MafA in the tissue. Most MafB-Sigma immunoreactivity in superficial laminae remains in the KOs suggesting that these cells strongly express a cross-reacting target and frequently do not express *mafb* (GFP negative). This is most likely c-Maf which is highly expressed in laminae I to III neurons.

**D,** As in **C,** for MafB-Novus antibodies. Unlike MafB-Sigma, there is little MafB-immunoreactivity in the het animal outside GFP+ cells reporting MafB expression; this includes superficial laminae cells. All immunoreactivity disappears in the KO animal.

**E,** Ventral horn co-localization of MafB-GFP and MafB-immunoreactivity obtained with Sigma and Novus antibodies. Graphs show the percentage of immunoreactive cells that are GFP+ (left) and the percentage of GFP+ cells that co-localize the indicated MafB antibody immunoreactivity (right). In one spinal cord ventral horn section from a het mouse (blue bars), we sampled 441 cells with MafB-Sigma immunoreactivity, but the same section had fewer MafB-GFP cells (n = 392) such that only a small number of cells co-localized both markers (n = 139). In total, 31.5% of MafB-Sigma immunoreactive cells expressed GFP and 35.5% of GFP+ cells had MafB-Sigma immunoreactivity. Conversely, in a serial section immunolabeled with the Novus antibody we detected fewer MafB-Novus immunoreactive cells (n = 149) than MafB-GFP cells (n = 215) and the large majority were GFP+ (92.0%, n = 137). There was almost no MafB-Novus immunoreactivity outside GFP+ cells. Thus, the MafB-Novus antibody is more restricted to GFP+ cells than the MafB-Sigma antibody. In addition, 63.7% of GFP+ cells express MafB-Novus immunoreactivity and while putative dorsal-MafB V1 cells express strong MafB-Novus immunoreactivity this is weak in Renshaw cells. In one MafB KO section (red bars) we detected 217 MafB-Sigma immunoreactive cells and 296 GFP+ cells with 39 cells co-localizing both markers. This corresponded to 17.9% of MafB-immunoreactive cells expressing GFP and 13.2% of the GFP+ cells expressing MafB-Sigma immunoreactivity. The MafB-Novus antibody showed no MafB-immunoreactivity in KO tissue sections with similar numbers of GFP+ cells (n = 241).

In summary, the MafB-Novus antibody detects MafB in the spinal cord very specifically. In contrast, the MafB-Sigma antibody strongly labels MafB-expressing cells along with some other cross-reacting proteins that may include MafA and c-Maf. Many superficial laminae interneurons may express the MafB-Sigma cross-reacting species: they are not detected with MafB-Novus and do not disappear in the MafB KO mouse. Renshaw cells and cells in the location of dorsal MafB-V1 cells are immunoreactive to both MafB antibodies, but detectability of Renshaw cells is better with the Sigma antibody (possibly because of additional cross-detection of MafA).

**F,** Western blots of MafA (Novus), MafB-Novus, and MafB-Sigma on nuclear-extracts from one WT (*mafb*$^{+/+}$), *one mafb* heterozygous (het, *mafb*$^{GFP/+}$) and one *mafb* knockout (KO, *mafb*$^{GFP/GFP}$), all E16 littermates. The same western blot was stripped and re-probed three times with the antibodies in the following order: MafB-Novus, MafA-Novus, and finally MafB-Sigma. Molecular weight standards are the same as in Supplementary Figure 1. The MafA antibody shows a single band just below the 37 kDa marker, aligning with a predicted molecular weight of 37.6 kDa. The band is of similar size in all three lanes suggesting no compensatory change in MafA expression in *mafb* hets and KOs. Neither of the MafB antibodies detected the MafA band in Western Blots, suggesting that any possible cross-reaction in tissue is due to IgG species detecting secondary or tertiary protein structures. MafB antibodies generated a double band, with the upper band being weaker than the lower band for MafB-Sigma compared to MafB-Novus. The immunoreactivity of the lower band to MafB-Sigma diminished in hets compared to WTs and diminished further in the KO. This band completely disappeared in the KO probed with MafB-Novus antibodies. This suggest that this band corresponds to MafB and occurs at approximately 40 kDa, slightly over the predicted 35.8 kDa molecular weight. Both antibodies detected a higher molecular weight band that does not change with gene dose. This could correspond to c-Maf with a larger predicted molecular weight, 38.5 kDa. Therefore, we performed a new western blot using a c-Maf antibody from Novus, and we found a correspondence between the upper band detected by both MafB antibodies with one of the bands in the c-Maf western blots. This suggests that in Western blots both MafB antibodies cross reacted with c-Maf.

Hu J, Huang T, Li T, Guo Z, Cheng L (2012) c-Maf is required for the development of dorsal horn laminae III/IV neurons and mechanoreceptive DRG axon projections. J Neurosci 32:5362-5373.

Frezel N, Ranucci M, Foster E, Wende H, Pelczar P, Mendes R, Ganley RP, Werynska K, d'Aquin S, Beccarini C, Birchmeier C, Zeilhofer HU, Wildner H (2023) c-Maf-positive spinal cord neurons are critical elements of a dorsal horn circuit for mechanical hypersensitivity in neuropathy. Cell Rep 42:112295.

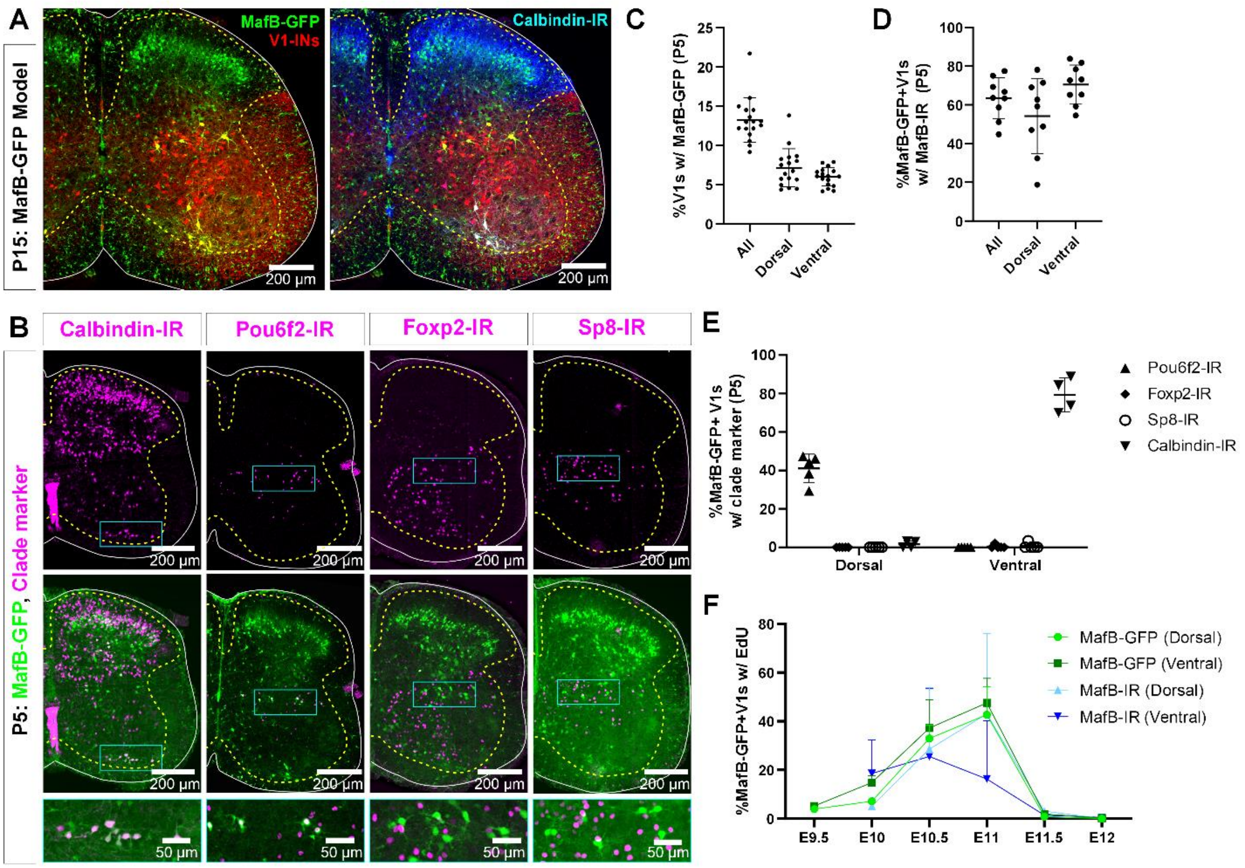


**Supplementary Figure 3: MafB-V1s visualized in a MafB-GFP mouse model.**

**A,** MafB-GFP+ cells in mature (P15) mice. Most MafB-GFP+ cells in lamina VII belong to the V1 lineage (tdTomato). They are found in the most dorsal and ventral regions of the distribution of V1s. Those that are also calbindin-IR fall in the distinctive ventral region occupied by Renshaw cells. In addition, there are many dorsal horn non-V1 MafB-GFP+ neurons. The small cells throughout the white and gray matter are microglia.

**B,** Expression of TFs and calbindin in MafB-GFP+ cells in neonatal (P5) mice. Top row, immunoreactivity for calbindin (Renshaw cells) and clade-specific TFs. Bottom row, superimposition with MafB-GFP (V1-tdTomato is not shown for simplicity). Compared to P15, a few more ventral horn neurons express MafB-GFP at P5, including some motoneurons, but MafB labeling is weaker in these cells. Within MafB-GFP+ V1 neurons, the two groups located at the most dorsal and most ventral regions correspond to the V1 neurons that retain MafB-GFP at P15 (see **A**). These groups are indicated with rectangles in the figure. The ventral group expresses calbindin. The dorsal group expresses Pou6f2 at this age. Little-to-no Foxp2-IR or Sp8-IR is found in either dorsal or ventral groups of MafB-GFP+ V1s.

**C,** Quantitation of MafB-V1 neurons in P5 mice. Around 13% of all V1s express MafB-GFP+, and the percentages located in the Renshaw cell area (“ventral”), or dorsal lamina VII (“dorsal”) are evenly split (n = 17 mice, 4 ventral horns each. Bars show SD).

**D,** More than half of the MafB-GFP+ V1 cells have detectable levels of MafB (Sigma) immunoreactivity at P5, in both the ventral and dorsal groups (n = 9 mice, 4 ventral horns each. Bars show SD).

**E,** V1- clade marker expression in dorsal and ventral P5 MafB-GFP V1 neurons. Dorsal MafB-GFP-V1s express Pou6f2, but do not express calbindin, Sp8 or Foxp2. Ventral MafB-GFP-V1s express calbindin (Renshaw cells) and do not express Pou6f2, Foxp2 or Sp8. (n=4-5 mice, 4 ventral horns each. Bars show SD).

**F,** EdU birthdating reveals similar proportions of EdU+ neurons at P5 for dorsal MafB-GFP vs MafB-IR neurons pulse-labeled at each embryonic timepoint (n=2 mice per timepoint per condition, 4 ventral horns each. Bars show SD). The mismatch in birthdates between MafB genetic and antibody labeling at E11 in the ventral group could arise because some E11-born ventral MafB-GFP V1s quickly downregulate MafB to levels that are undetectable with antibodies.

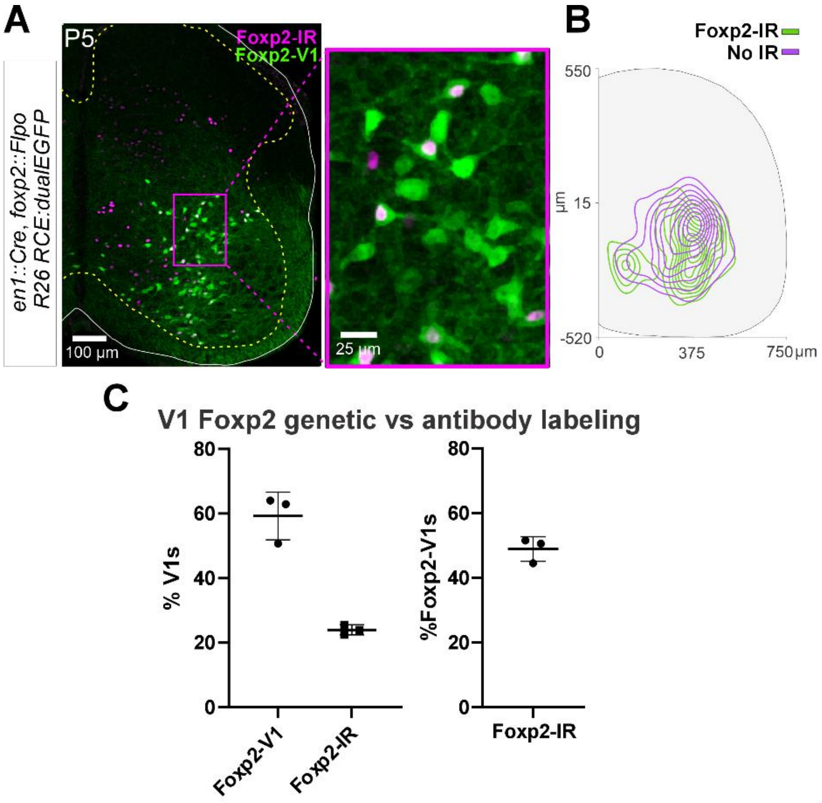


**Supplementary Figure 4. Postnatal downregulation of Foxp2 expression in Foxp2-V1 cells.**

**A,** Confocal image of *en1::Cre*, *foxp2::Flpo,* R26::RCE:dualEGFP genetic labeling (green) immunostained for Foxp2 protein expression (magenta). Low magnification 2D projection at the left. Higher magnification of the indicated area at the right. Only a proportion of lineage label Foxp2-V1s express Foxp2 protein by P5.

**B,** Density contours demonstrate that lineage labeled Foxp2-V1s with or without Foxp2 protein expression occupy similar areas in the spinal cord at P5 (n=3 mice, 3 ventral horns per animal).

**C,** <u>Left</u>, Foxp2-V1s make up roughly 60% of all V1s when measured with genetic labeling, but only about 30% of V1s maintain detectable Foxp2 expression at P5 (n is same as B; error bars show SD).
<u>Right</u>, around half of Foxp2-V1s maintain detectable levels of Foxp2 at P5 (same data).

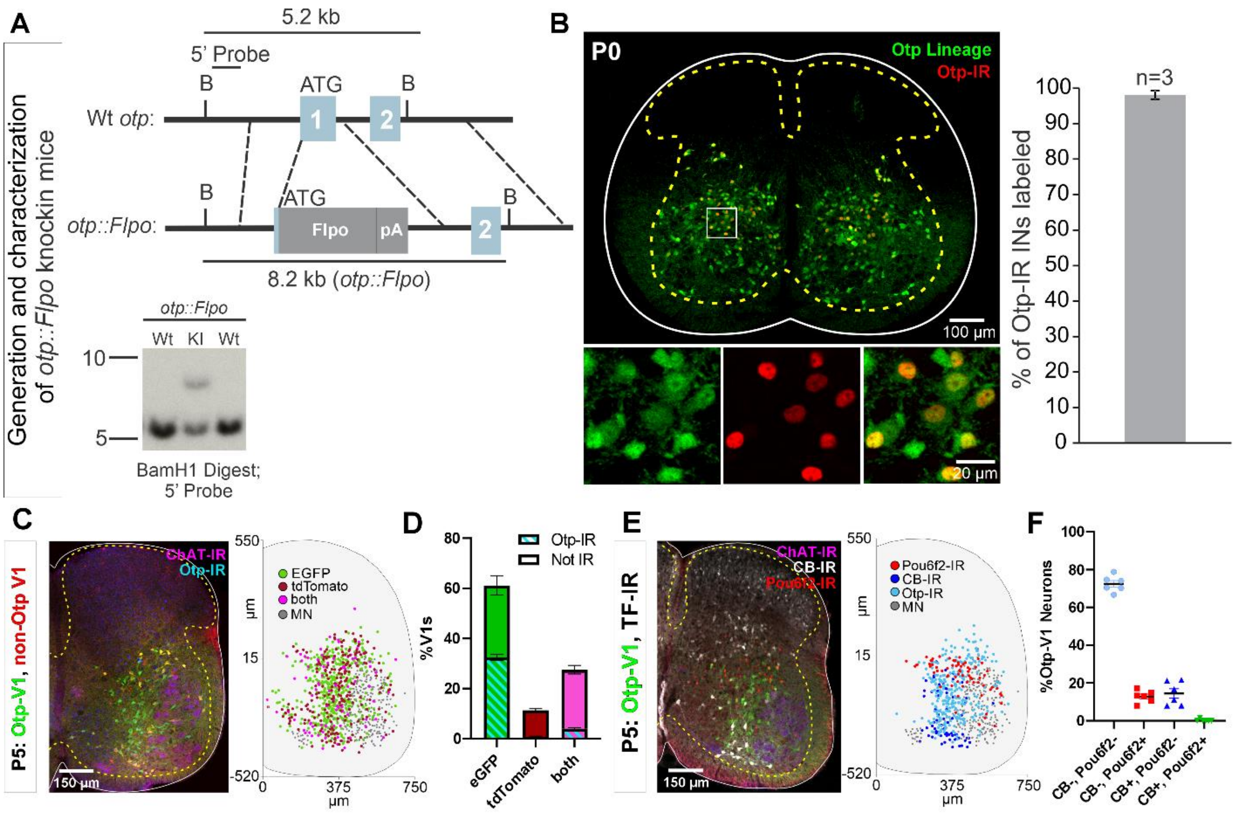


**Supplementary Figure 5. Generation of Otp-Flpo mice and intersection with En1-cre mice to target Otp-V1 cells.**

**A,** Targeting strategy to generate o*tp::flpo* mice. Flpo was inserted into the ATG in the 1st exon of the Otp genomic locus. Dotted lines represent approximate regions of homology in the targeting vector. Southern blot (bottom) of BamH1-digested genomic DNA with a 5' probe external to the targeting vector identifies a 5.2 kb wild-type fragment, and an 8.2 kb knock-in fragment. Not shown: Deletion of selectable neomycin resistance gene flanked by loxP sites by crossing to Protamine-Cre mice, which recombines the floxed PGK-Neo cassette in the male germline.

**B,** Left, P0 lumbar spinal cord of o*tp::flpo*, RCE.fsf.GFP mice, demonstrating expression in Otp-IR interneurons. Right, 98.1 ±1.2% (mean ± SEM, n = 3 mice) of Otp-expressing cells are labeled by the reporter.

**C,** Intersection of *otp::flpo* and *en1::cre* using the dual color strategy with simultaneous expression of the Ai9 tdT and RCE-DC eGFP reporters. V1 cells that express *otp* are labeled with EGFP (green). In these cells the Ai9-tdT reporter is effectively deleted by Flpo recombination dependent on the level of *otp* expression: strong (EGFP only) and weak (EGFP and tdT). V1 cells that express tdT only (red) never express *otp*. In addition, the sections were labeled with antibodies for Otp (light blue) to reveal cells that retained expression of Otp at P5 and ChAT (deep blue) to localize the motor pools. Most V1s express EGFP, with tdT-only cells being a minority. Right, cell plot positions of some of the cell types identified in these sections (one mouse 6 ventral horns in L4/L5). Most cells are Otp-V1s and are shown here as green (eGFP only) and pink dots (EGFP and tdT).

**D,** Quantification of cells with EGFP only (green), tdT only (red), or both (pink) (n=12 ventral horns from 2 mice, error bars show SD). 88.6% of V1s (1,278 V1 cells analyzed in total) expressed EGFP; 60.8% were EGFP only, 27.8% were "yellow" and surprisingly only 11.4% were tdT only. As expected, P5 Otp expression detected with antibodies was absent in most tdT labeled V1 cells (93.5% of cells) and "yellow" tdT+eGFP V1 cells (85.4%), but also in a significant proportion of eGFP labeled V1 cells (46.7%).

**E,** V1-cells transiently expressing Otp in embryo included cells of other clades. This was examined in $en1^{cre/+}$::$otp^{flpo/+}$::*RCE:dual-eGFP* mice. Detection of Pou6f2, calbindin and Otp in genetically labeled Otp-V1 cells at P5. Left confocal image and right cell plot (n=6 ventral horns from 1 mouse). Pou6f2-Otp-V1 cells concentrate in a dorsal band within the ventral horn. Calbindin-IR Otp1-V1 cells concentrate in the Renshaw region (ventral most region) and others are in the dorsal region of the ventral horn. Otp-V1 cells retaining Otp expression at P5 occupy all dorsoventral positions in the lateral spinal cord.

**F,** Pou6f2 or calbindin immunoreactivity (-IR) in Otp-V1 cells (n=397) in 6 sections (each dot) in L4/L5 from one animal. Pou6f2 was detected in 12.8% of Otp-V1 cells at P5. In addition, 14.4% of Otp-V1 cells were calbindin+ and this included many in the Renshaw cell ventral region and few others located more dorsally. One rare dorsal Otp-V1 cell contained both Pou6f2 and calbindin (included in both percentages above). By limiting the analysis to ventral Otp-V1 interneurons in the Renshaw area we estimated that 8.3% of them are Renshaw cells.

In conclusion, the Otp-V1 lineage includes cells from several V1 clades. Many downregulate Otp expression before birth, thus at P5 all Otp-expressing cells are restricted to the Foxp2-V1 lineage. In addition, many medial non-V1 Foxp2 cells also express Otp. Therefore, to specifically target the lateral group of proprioceptive Otp-Foxp2-V1 cells tightly associated to the LMC, a triple genetic intersection or alternatively, postnatally timed Otp-dependent recombination is necessary.

TA CTB-labeling and LG RV-mCherry monosynaptic tracing

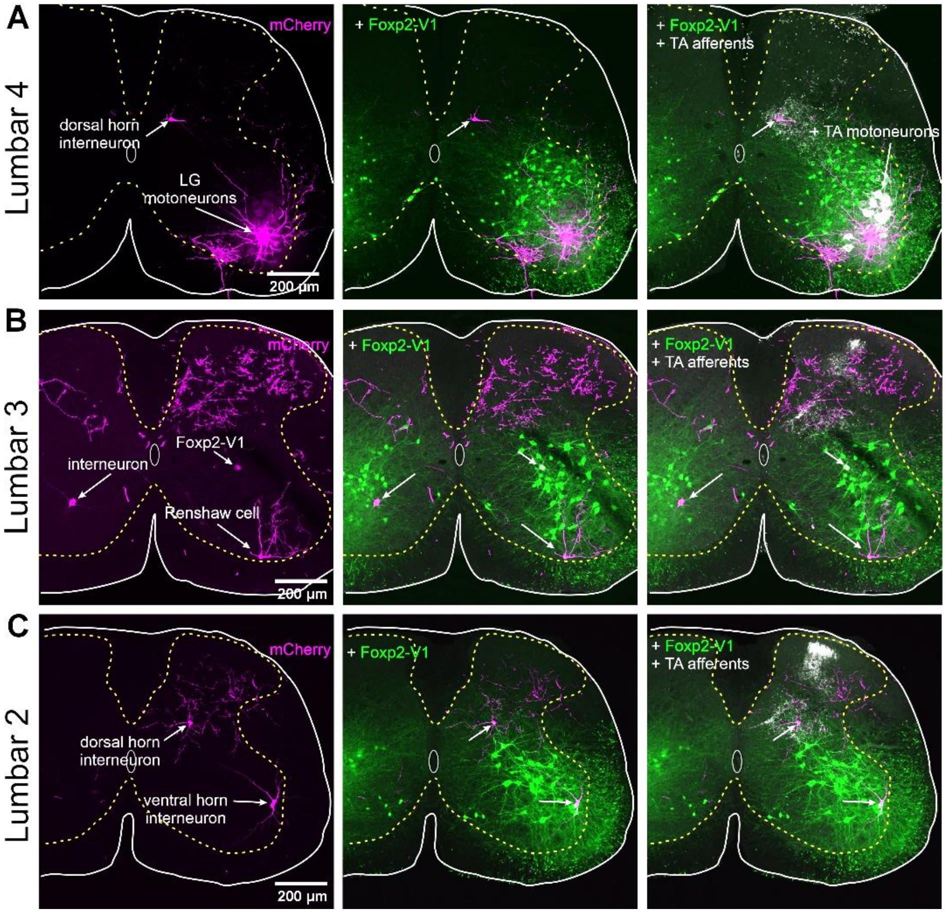


**Supplementary Figure 6. Labeling of motoneurons and interneurons, but only limited labeling of primary afferents with RVΔG-mCherry.**

All confocal images are serial to those shown in Figure 8: AAV1-G was injected at P4 and RVΔG-mCherry in LG and CTB in TA at P15, the animal sacrificed at P22, 7 days after RV injection.

**A,** Left panel. Lumbar 4 section showing RVΔG-mCherry labeling in LG motoneurons and one dorsal horn interneuron in medial LV. Many dendrites, motor axons and collaterals located in the Renshaw cell area are labeled. There are no mCherry labeled primary afferents in the dorsal root, dorsal columns or dorsal horn in this segment (the same for Lumbar 5).

Middle panel. Added Foxp2-V2 lineage labeling (EGFP). The dorsal horn interneuron is far from the ventral horn region containing Foxp2-V1 interneurons.

Right panel. Added CTB labeling injected in the TA. TA motoneurons and TA primary afferents are respectively retrogradely and anterogradely labeled. TA afferents concentrate in medial lamina V and contact the dorsal horn mCherry interneuron monosynaptically connected to the LG motor pool.

**B,** Left panel. Lumbar 3 section showing extensive RVΔG-mCherry axon labeling in the dorsal horn and three different types of ventral horn interneurons monosynaptically connected to the LG motor pool. This lumbar segment contained the denser dorsal horn axon labeling found in this animal. Some of these axons may arise from primary afferents since they can be traced to the dorsal column. These axonal arbors are far from ventral horn interneurons and have no intermingled mCherry-labeled interneurons. The three mCherry lumbar interneurons labeled in this section are located in different ventral horn positions: contralateral LVII, ipsilateral Renshaw cell area and ipsilateral middle of LVII.

Middle panel. Added Foxp2-V2 lineage labeling (EGFP). Interneurons located in the contralateral spinal cord and in the ipsilateral Renshaw cell area are not Foxp2-V1. The cell in the middle of LVII is a Foxp2-V1 interneuron.

Right panel. Added CTB labeling injected in the TA. Only TA afferents are visible in this segment. They occupy medial LV as in Lumbar 4 (**A**). TA afferents also project to the superficial laminae in this segment.

**C,** Left panel. Lumbar 2 section showing RVΔG-mCherry labeling of dorsal and ventral horn interneurons. This lumbar segment shows some axon labeling in the dorsal horn. Some may arise from primary afferents.

Middle panel. Added Foxp2-V2 lineage labeling (EGFP). mCherry-labeled interneurons in this section are not Foxp2-V1 interneurons.

Right panel. Added CTB labeling injected in the TA. Only TA afferents are visible in this segment. Similar to the lumbar 3 segment (**B**), TA afferents project to medial lamina V and also to superficial laminae. TA afferents contact the dorsal horn mCherry interneuron in medial lamina V.

Detailed examination of serial sections from L2 to L6 in this animal and others revealed limited RV labeling of primary afferents. When these are present, they are restricted the dorsal horn of lumbar 3 and 2 segments. This suggests that there is only limited infection of sensory afferents when animals are intramuscularly injected with this strain of virus (B16 RVΔG) at P15. Therefore, mCherry ventral horn interneurons are most likely transsynaptically labeled in the retrograde direction from starter motoneurons infected with both RVΔG-mCherry and AAV1-G that were injected in the muscle.
